# Efficacy of the first bioactive peptide from the pearl

**DOI:** 10.1101/2023.10.10.561660

**Authors:** Chaoyi Wu, Zehui Yin, Yayu Wang, Xinjiani Chen, Bailei Li, Qin Wang, Liping Yao, Zhen Zhang, Xiaojun Liu, Rongqing Zhang

**Affiliations:** Key Laboratory of Freshwater Aquatic Genetic Resources, Shanghai Ocean University, Ministry of Agriculture, Shanghai 201306, China; Department of Biotechnology and Biomedicine, Yangtze Delta Region Institute of Tsinghua University, Zhejiang 314000, China; Taizhou Innovation Center, Yangtze Delta Region Institute of Tsinghua University,Zhejiang 318000,China; Zhejiang Provincial Key Laboratory of Applied Enzymology, Yangtze Delta Region Institute of Tsinghua University, 705 Yatai Road, Jiaxing 314006, China

**Keywords:** Angiotensin-1-converting enzyme (ACE)-inhibitory peptide, inhibition kinetic, molecular docking, pearl matrix proteins

## Abstract

Pearls have high medicinal value. In the present study, we discovered the first bioactive peptide in pearls. The bioactive peptide, KKCHFWPFPW, was a novel angiotensin I-converting enzyme (ACE)-inhibitory peptide derived from the pearl matrix of *Pinctada fucata*. It was screened and identified using quadrupole time-of-flight mass spectrometry. The molecular weight of the peptide was 1417.5 Da, and its theoretical isoelectric point was 9.31. The half-maximal inhibitory concentration of the peptide was 4.17μM, as determined by high-performance liquid chromatography. The Lineweaver–Burk plot showed that this peptide competitively inhibited ACE activity. As the peptide concentration increased, the ACE inhibition rate also increased. The molecular docking was simulated using Maestro 2022-1 Glide software to understand the potential mechanisms underlying the ACE-inhibitory activity of KKCHFWPFPW. These results indicated that the peptide from the *P. martensii* pearl matrix might be a potential source of antihypertensive peptides.

## 1. Introduction

Hypertension is one of the main diseases threatening human health. It is characterized by the contraction of the dynamic pulse or the rise of diastolic pressure. It is highly correlated with heart, brain, and blood vascular diseases. Hypertension is an independent risk factor for cardiovascular diseases, which are frequent, chronic, and age-related [1]. Therefore, antihypertensive treatment can prevent cardiovascular diseases.

Angiotensin I-converting enzyme (ACE) is an important enzyme in the renin-angiotensin system, which is an essential hormone system responsible for the homeostasis of blood pressure in mammals. ACE removes the C-terminal dipeptide from its substrate precursor peptide angiotensin I to produce angiotensin II, which is a potent vasoconstrictor that blocks the vasodilating properties of bradykinin, ultimately causing an increase in blood pressure [2]. These two functions make ACE an ideal target for treating hypertension, heart failure, type 2 diabetes mellitus, and diabetic nephropathy.

The first ACE inhibitors were isolated from the venom of the snake *Bothrops jararaca* [3]. Subsequently, ACE inhibitors such as captopril, lisinopril, enalapril, and fosinopril, which are commonly used to treat hypertension, were developed [4]. However, these synthetic ACE inhibitors may have adverse effects such as cough, allergic reactions, taste disorders, and skin rashes [5]. Therefore, identifying bioactive peptides that can inhibit ACE, have no toxicity, can lower blood pressure, and have no hypotensive effect on people with normal blood pressure is extremely important. Bioactive peptides are also involved in immunoregulation and are easy to digest and absorb. Although the activity of bioactive ACE-inhibitory peptides is lower than that of chemosynthetic ACE inhibitors, the bioactive peptides are safer, low-priced, and a source of essential amino acids. Therefore, bioactive ACE-inhibitory peptides have traits superior to those of chemosynthetic ACE inhibitors [6, 7].

The ocean is a treasure trove of biological resources containing abundant bioactive ingredients, such as active peptides, active polysaccharides, and polyunsaturated fatty acids. Among these, bioactive peptides extracted from fish, shellfish, seaweed, and other marine organisms have antioxidant, blood pressure–regulatory, antithrombosis, and immunoregulatory functions. With an increase in the exploitation and utilization of marine biological resources, identifying ACE inhibitors from marine-derived proteins has become promising.

The bivalve *Pinctada martensii*, also known as *Pinctada fucata*, is an essential mariculture species and the main mother shellfish for pearl production. Additionally, modern medical research has shown that natural seawater pearls can be used to improve human immunity, protect against senility, whiten skin, and provide calcium. Therefore, these pearls have a large potential market in the fields of medicine, dietary supplementation, and cosmetic production.

Pearls are formed via mineralization in the shells of bivalve mollusks. The main component of the shell is calcium carbonate (∼95%), and the remaining components are a protein and chitin organic matrix (∼5%). The organic matrix includes organic macromolecules such as proteins, polysaccharides, and lipids. The protein components are collectively referred to as matrix proteins. They are mainly involved in constructing the organic framework and regulating calcium carbonate nucleation and crystallization [8]. Although the organic matrix proportion is small, it controls the whole shell formation process. Matrix proteins precisely regulate crystal nucleation, morphology, growth, and orientation [9, 10]. The nacre of mollusks has a structure similar to that of pearls; the shell matrix proteins from the nacre are considered essential mediators of pearl formation [11, 12].

The ACE inhibitor assessed in this study was a pearl matrix peptide from *P. fucata*. Although pearls have been studied extensively, no antihypertensive peptide was derived from them prior to this study. We used the classic Lineweaver–Burk model to investigate the ACE-inhibitory activity of pearl matrix peptide KKCHFWPFPW *in vitro* and explore the mode and mechanism of inhibition of potential inhibitory proteins. The results of this study provided theoretical and experimental bases for using a pearl matrix active peptide as a new bioactive component.

## 2. Materials and Methods

### 2.1 Materials and chemicals

Nucleus-free pearls from *P. fucata* were purchased from Kanjiang, Guangdong Province, China. ACE, hippuryl-L-histidyl-L-leucine (HHL), hippuric acid (HA), trifluoroacetic acid (TFA), boric acid, sodium chloride (NaCl), acetonitrile, and hydrochloric acid (HCL) were obtained from Sigma–Aldrich (MO, USA). All chemicals used in this study were of analytical grade.

### 2.2 Extraction and enzymolysis of pearl matrix proteins

The pearls were cleaned and crushed with a shredder. Next, 1.5M EDTA was added to the collected pearl powder and stirred for 24 h at 4°C to decalcify the powder. The supernatant was collected by centrifugation at 11962 g and filtered through a 0.2-μm microporous filter membrane. The filtrate was placed in a chromatography freezer at 4°C for low-temperature dialysis for 48 h. The pearl matrix proteins were obtained by freezing and drying the filtrate.

Trypsin (at 5% of the pearl matrix proteins with activity of 10,000 U/g) was added to the pearl matrix proteins for enzymatic hydrolysis. Digestion was performed at pH 8.0. The trypsin was hydrolyzed for 2 h at 50°C and then inactivated in a boiling water bath. The supernatant was collected by centrifugation at 11962 g. The filtrate < 3 kDa was collected after ultrafiltration. Enzyme-digested pearl matrix proteins were obtained after concentration and freeze-drying.

### 2.3 Separation and purification of pearl matrix proteins with ACE-inhibitory activity

High-performance liquid chromatography (HPLC) was used to purify the enzyme-digested pearl matrix proteins. In a C18 column, the mobile phase A was deionized water containing 0.1% TFA, and the mobile phase B was acetonitrile containing 0.1% TFA. The ultraviolet detection wavelength was 280 nm, the flow rate was 1 mL/min, and the retention time was 14 min. The pearl matrix proteins with ACE-inhibitory function were obtained by concentration and freeze-drying.

### 2.4 Assay of ACE-inhibitory activity

For this assay, the reaction system consisted of 5mM HHL, the pearl matrix protein inhibitor, and 200 U/L of ACE in 0.1M sodium borate buffer (pH 8.3, 0.3M NaCl). Six inhibitor concentrations were tested (5, 10, 20, 30, 40, and 50 μL, supplemented with 0.1M borate buffer to 180 μL). The control group contained 0.1M borate buffer without inhibitor. First, HHL and the inhibitor were mixed and preheated for 5 min at 37°C. Next, 20 μL of ACE was added, and the mixture was kept in a shaking water bath for 30 min at 1.72 g and 37°C. Finally, 0.2 mL of 1 mol/L HCL was added to terminate the reaction. The concentrations of HHL and its hydrolysate HA were determined using an HPLC on a C18 column. The mixture was analyzed using isometric elution for 25 min in 75% mobile phase A (pure acetonitrile) and 25% mobile phase B (deionized water containing 0.5% TFA) at a flow rate of 0.5 mL/min. The ultraviolet detection wavelength was set to 228 nm. ACE can stably degrade HHL and generate HA. Hence, once the activity of ACE is inhibited, the amount of HA produced in the sample also decreases. The ACE-inhibitory activity of the substrate was calculated using the following formula:

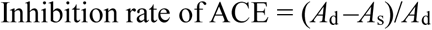

where *A*_s_ is the detection peak area of HA in the sample group and *A*_d_ is the peak area of HA detection in the blank group. From the results, we calculated the half-maximal inhibitory concentration (IC_50_) value.

### 2.5 Determination of molecular weight and amino acid sequence

The molecular weight and amino acid sequence of the purified peptides from pearl matrix proteins were determined using a quadrupole time-of-flight mass spectrometer (Q-TOF MS; Micromass, Cheshire, UK) coupled with an electrospray ionization (ESI) source. The purified peptide dissolved in methanol/water (1:1, *v*/*v*) was infused into the ESI source, and the molecular weight was determined by doubly charged (M+2H)2+ state analysis in the mass spectrum. Following the molecular weight determination, the peptide was automatically selected for fragmentation, and the sequence information was obtained by tandem MS analysis.

### 2.6 Determination of ACE-inhibitory kinetics

The ACE inhibition pattern of peptide KKCHFWPFPW was analyzed using the Lineweaver–Burk diagram of 1/*V* and 1/HHL. Various HHL concentrations (0.5, 1, and 2 mM) and peptide concentrations (0, 17.63, 35.26, 70.52, 105.78, 141.04, and 176.3 μM) were incubated with the ACE solution. The Michaelis constant (*K*_m_) and maximum velocity (*V*_max_), along with the inhibition type of the peptide, were determined graphically using Lineweaver–Burk plots. *V*_max_ and *K*_m_ were calculated as the *Y*-intercept and *X*-intercept of the main curve, respectively. All experiments were conducted in triplicate.

### 2.7 Molecular docking

The molecular docking of the pearl peptide with ACE was performed using Maestro 2022-1 Glide. The crystal structure of ACE (ProteinData Bank ID: 1O8A) was downloaded from the PDB database. The structure and minimum energy forms of the target peptide were calculated automatically according to the amino acid sequence of the target peptide before molecular docking. The type of interaction between the docking receptor and the ligand was analyzed.

## 3. Results and Discussion

### 3.1. Identification of peptides from pearl matrix proteins

In the present study, UPLC-Q-TOF-MS/MS and UniProt search were used to analyze and detect the fragment spectra so as to identify the peptides in the pearl matrix proteins.

Among the peptides obtained by enzymolysis from pearl matrix proteins, one peptide, KKCHFWPFPW, was predicted to be an ACE-inhibitory peptide according to the affinity analysis by molecular docking. Therefore, the KKCHFWPFPW was purified and isolated to clarify the mechanism of ACE-inhibitory activity. The molecular weight of the ACE inhibitor (ACEI) was 1417.5 Da, as determined by MS/MS (Fig. 1a). Also, its amino acid sequence was Lys-Lys-Cys-His-Phe-Trp-Pro-Phe-Pro-Trp (Fig. 1b). The theoretical isoelectric point was 9.31, and the total average hydrophilicity was −0.790. It was an alkaline hydrophilic protein.

**Figure 1.**
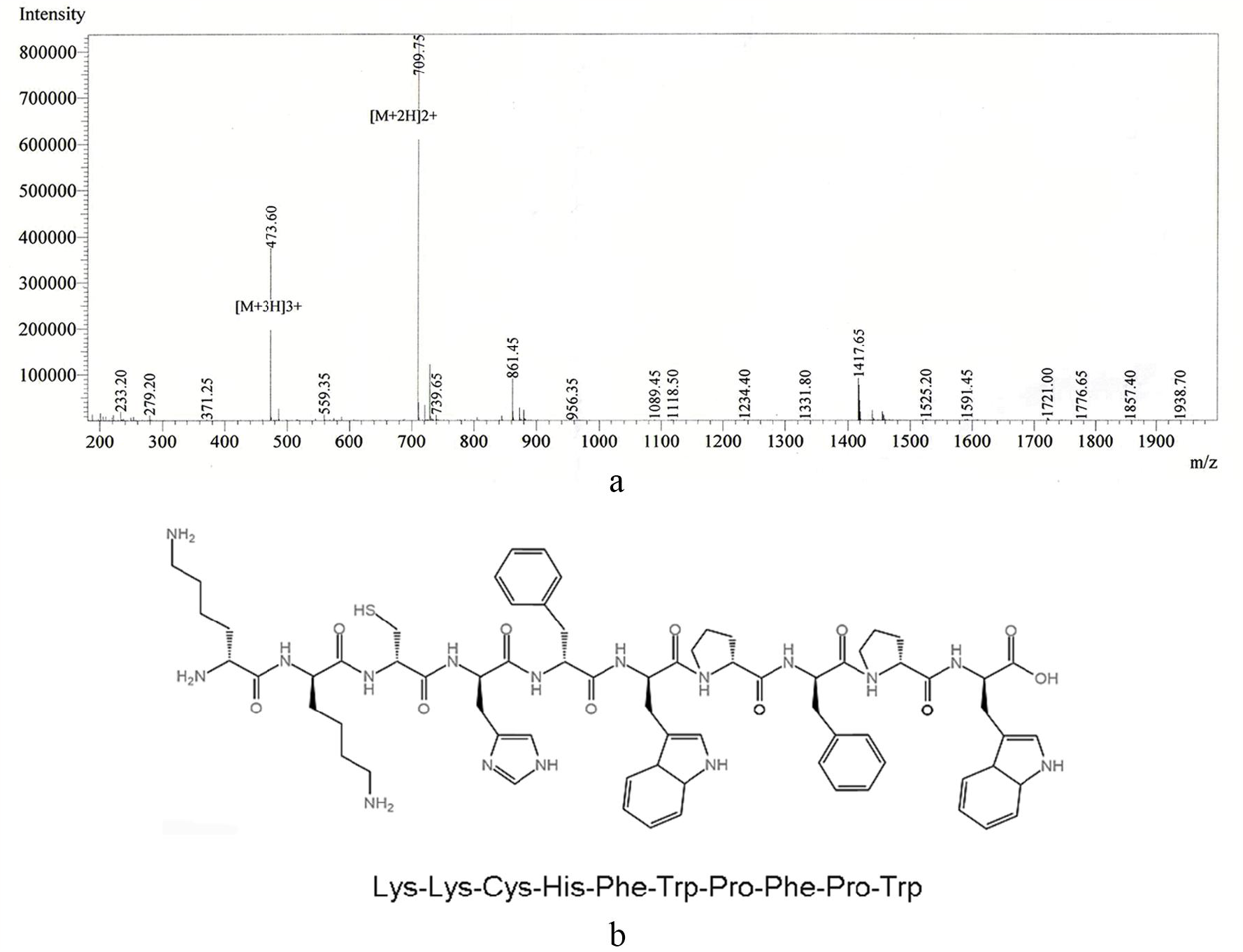
Identification of KKCHFWPFPW by (a) MS and (b) UPLC-Q-TOF-MS/MS.

The present study identified the peptide KKCHFWPFPW from the pearl matrix protein hydrolyzed by trypsin for 2 h at 50°C and pH 8.0. This was the first attempt to extract ACE-inhibitory peptides from mollusk shell pearls. The result of UniProt search showed that the peptide KKCHFWPFPW could be matched with the peptides in two kinds of *Pinctada fucata* matrix proteins, which were named as lysine-rich matrix protein 6 and lysine-rich matrix protein 9. The location of KKCHFWPFPW (marked in red) in lysine-rich matrix protein 6 and lysine-rich matrix protein 9 from the UniProt database is shown in Figures 2 and 3, respectively.

**Figure 2.**
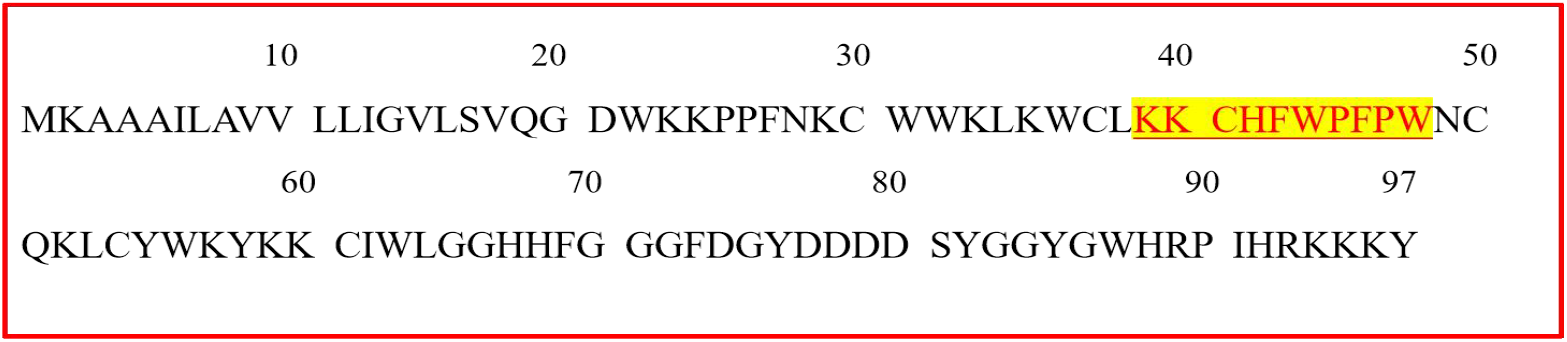
Location of KKCHFWPFPW in lysine-rich matrix protein 6 (KP940480).

**Figure 3.**
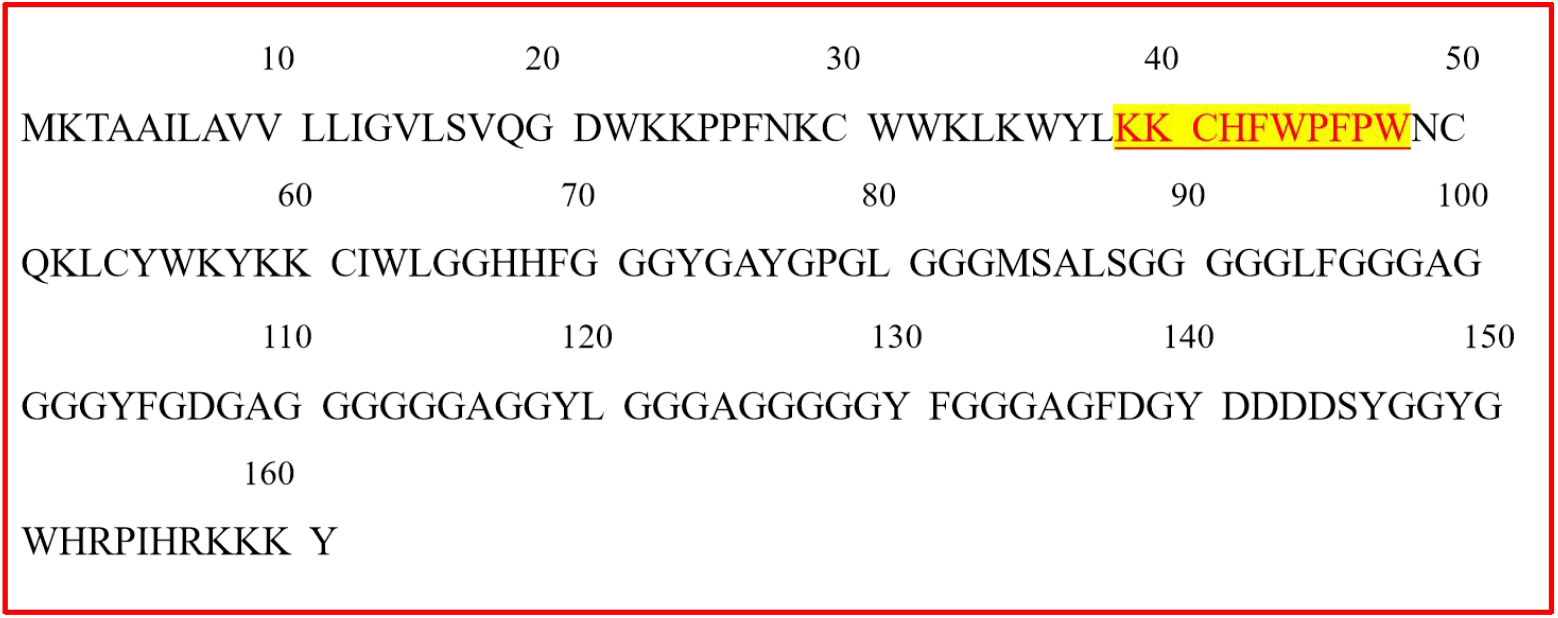
Location of KKCHFWPFPW in lysine-rich matrix protein 9 (KP940483).

Moreover, several studies have reported on the peptides of hydrolyzed proteins from various natural sources with ACE-inhibitory activity. The sources of ACE inhibitors are divided into four groups: plant sources, animal sources, microbial fermentation sources, and marine sources. The animal sources include milk, fish, and eggs. The first ACE inhibitors were isolated from the venom of *B. jararaca* in 1965 [3]. Subsequently, eggs, which are rich in proteins and many essential amino acids, were found to be an ideal raw material for ACE inhibitors. Examples include WESLSRLLG [11] from ostrich albumen and LKYAT and TNGIIR from egg white [12]. Milk is another vital source of ACE inhibitors because of its balanced nutrition and a wide variety of proteins [13].

Plant sources of ACE-inhibitory peptides include legumes [14], cereals [15], and seeds [16]. ACE inhibitors from plant sources have three to eight amino acid residues and special structures that make them more hydrophobic, conferring strong ACE-inhibitory activity. Microbial fermentation sources include *Lactobacillus* and other species. For example, *Lactobacillus plantarum* secretes extracellular protease to hydrolyze soybean proteins during the fermentation process, producing many ACE inhibitors [17]. Marine sources include fish [18], mollusks [19, 20], aquatic products [21, 22], and seaweed [23].

Many ACE-inhibitory peptides have been isolated and identified from mollusks, which have the potential for lowering blood pressure. For example, Liu et al. [24] isolated and identified VVCVPW and found that it bound to ACE via interactions with His383, His387, and Glu411 residues in *Mactra veneriformis*. Li et al. [25] studied the activity of ACE with razor clam (*Sinonovacula constricta*) hydrolysates using five proteases. Wang et al. [26] isolated and characterized a purified peptide (VVYPWTQRF) from oyster (*Crassostrea talienwhanensis*) protein hydrolysate. Wu et al. [27] used alcalase, followed by papain, to hydrolyze abalone (*Haliotis discus hannai*) gonads to produce ACE-inhibitory peptides. *P. martensii* tissues were hydrolyzed by protease, and ACE-inhibitory peptides were obtained after the separation and extraction of the hydrolysate [28].

Various studies showed that the amino acid composition, structure, and length of the antihypertensive peptide significantly affected ACE-inhibitory activity. Daskaya-Dikmen et al. [29] explored the structure–activity relationship of ACE-inhibitory peptides and reported that peptides with a rich hydrophobic amino acid content, such as Ala, Val, Leu, Ile, Phe, Pro, Trp, and Met, at the C-terminal residues had a strong effect on ACE binding. Kumar et al. [30] built a platform for predicting, screening, and designing antihypertensive peptides. They reported that residue Pro was highly abundant in antihypertensive peptides. Cushman and Cheung [31] reported that Trp, Tyr, Pro, and Phe at the C-terminal, as well as branched aliphatic amino acids at the N-terminal, were suitable for ACE binding as competitive inhibitors. Additionally, Norris et al. [7] found that shorter peptides might have stronger ACE-inhibitory activity. The molecular weight of KKCHFWPFPW was 1417.5 Da, and the number of amino acids was 10. Hence, the identified peptide was consistent with the basic characteristics of the aforementioned antihypertensive peptides. Based on the aforementioned conclusions, the sequence of pearl matrix proteins was found to contain Pro, Trp/Pro, and Phe, which helped improve its ACE-inhibitory ability. Moreover, the affinity of peptides to ACE protein was evaluated by molecular docking. KKCHFWPFPW had a relatively high affinity toward ACE, which was evaluated as −10.6691 using the Maestro 2022-1 Glide module. Therefore, the KKCHFWPFPW can be selected for further investigation.

### 3.2. ACE-inhibitory activity determination

Figure 4 depicts the HPLC chromatograms of the peptide KKCHFWPFPW at concentrations of 17.63, 35.26, 70.52, 105.78, 141.04, and 176.3 μM and a blank control group. The 11- to 12-min peak in the figure is HA, and the 20- to 21-min peak is the substrate HHL.

**Figure 4.**
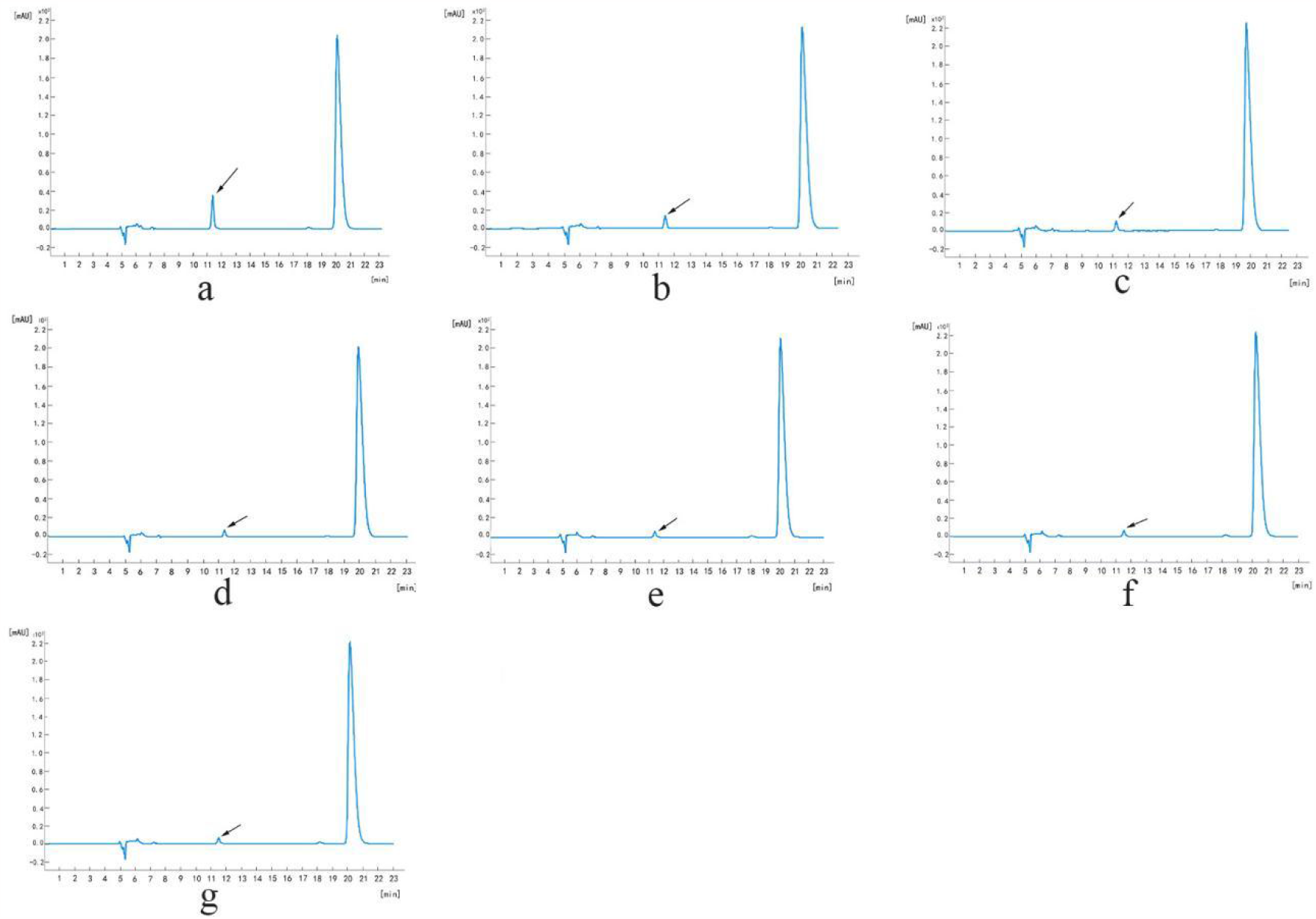
ACE inhibition of small-molecule active peptides by HPLC. Concentration of polypeptides: (a) 0 μM (control group); (b) 17.63 μM; (c) 35.26 μM; (d) 70.52 μM; (e) 105.78 μM; (f) 141.04 μM; and (g) 76.30 μM.

The ACE-inhibitory activity was calculated using the following formula: ACE inhibition rate = (*A*_s_ – *A*_d_)/*A*_d_ (*A*_s_ is the peak area of HA detection in the sample group, and *A*_d_ is the peak area of HA detection in the blank group). The IC50 value was determined using the regression equation: *Y* = −(4.7 × 10^−3^)*X*^2^ + 1.1352*X* + 18.908 (*R*^2^ = 0.8055).

The ACE inhibition rates were 53.2%, 65.9%, 74.4%, 77.2%, 78.1%, and 79.1%. The IC50 value of the peptide was 4.17μM. In the present study, the ACE-inhibitory activity of the peptide KKCHFWPFPW released by trypsin digestion was evaluated, as shown in Figure 5. Therefore, the peptide KKCHFWPFPW could be considered as a potential ACEI.

**Figure 5.**
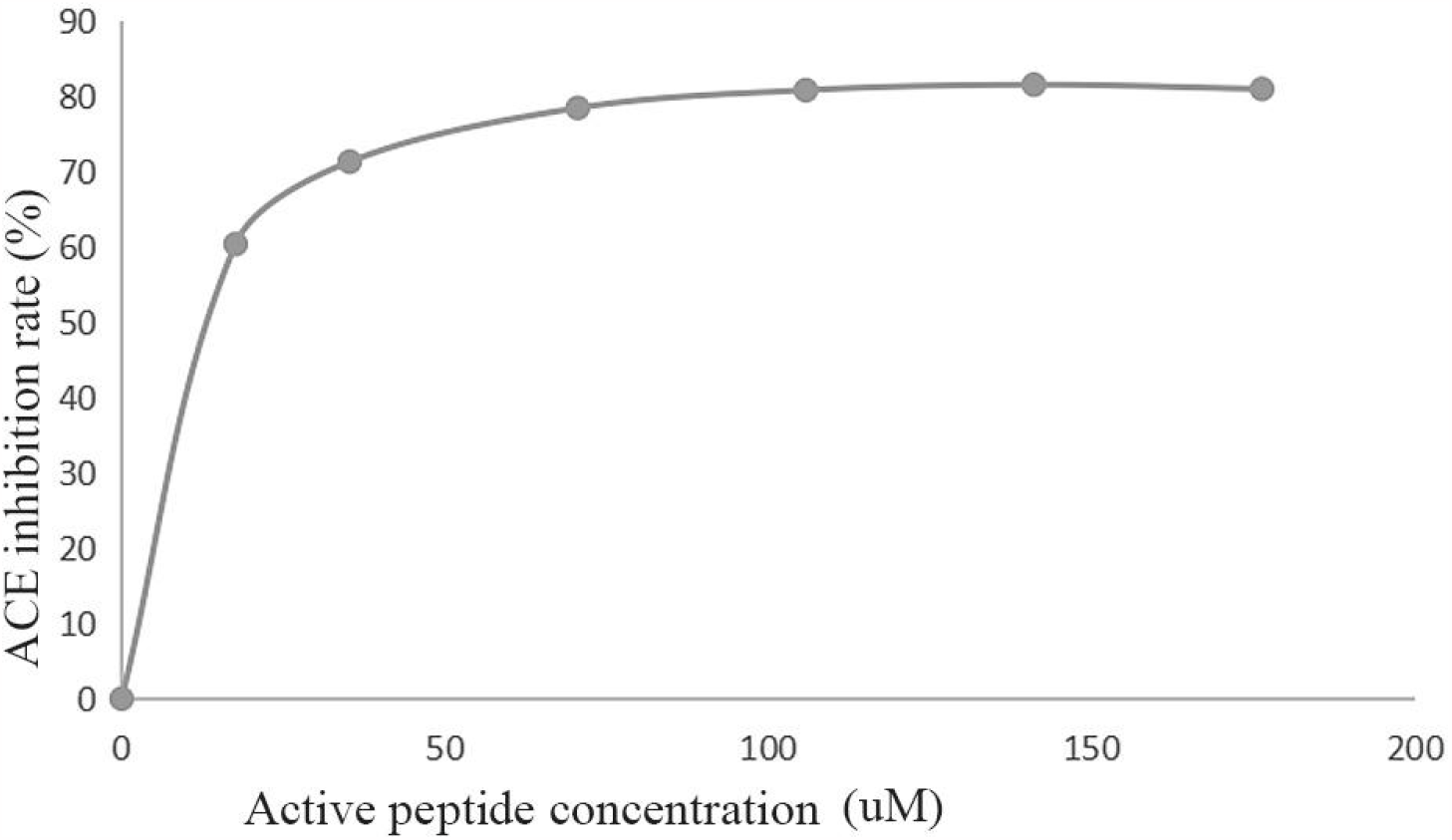
ACE inhibition activity diagram.

### 3.3. Inhibition pattern of the ACE-inhibitory peptide

The inhibition of ACE by ACE-inhibitory peptide was estimated using Lineweaver–Burk plot analysis to evaluate the mechanism of action of the peptide. The *V*_m_ values decreased with the increase in the pearl matrix peptide concentration (Fig. 6), confirming that the peptide likely blocked the substrate from binding to the ACE active site. When the ACEI peptide was added to the reaction, *K*_m_ values were higher than that of the control, indicating that a higher substrate concentration was essential for the ACE-catalyzed reaction. The regression curves of different concentrations of the pearl matrix peptide intersected at the 1/[*v*] axis, indicating that the inhibition mode was competitive inhibition.

**Figure 6.**
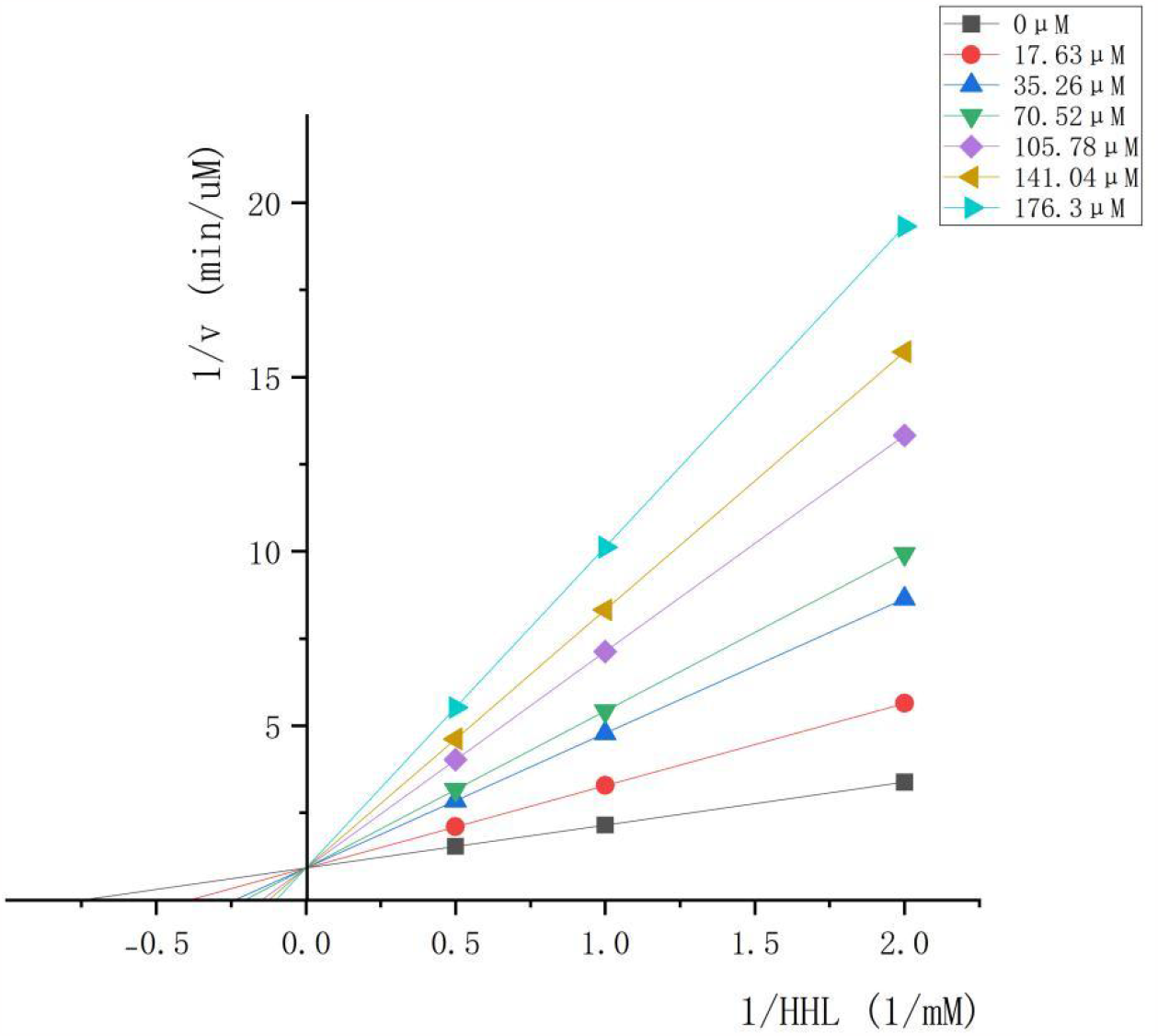
Lineweaver–Burk plot of ACE inhibition by the pearl matrix protein. 1/[HHL] and 1/[*v*] represent the reciprocal substrate concentration and velocity, respectively.

The ACE-inhibitory peptides are classified based on different inhibition mechanisms. They are divided into three groups: substrate-type peptide, prodrug-type peptide, and inhibitor-type peptide [32]. The activity of the substrate-type peptide is increased by pre-incubation with ACE. The prodrug-type peptide is converted by ACE or gastrointestinal enzymes into a true inhibitor with higher activity. The inhibitor-type peptide is a true inhibitor, and its activity does not change after treatment with ACE or other enzymes. The main inhibition models of ACE-inhibitory peptides are classified into competitive and noncompetitive inhibition. Competitive inhibition is the interaction of the inhibitor with active enzyme sites to prevent substrate binding [33]. Noncompetitive inhibition occurs when the inhibitor molecule has a binding affinity for free enzyme and enzyme–substrate complex [34].

According to the Lineweaver–Burk plot, the peptide KKCHFWPFPW identified in this study was a true inhibitor because it competed with the substrate HHL for the binding sites of ACE. Only a few peptides exhibited noncompetitive inhibition (i.e., the peptide did not compete with the substrate and only bound to the enzyme – substrate complex). Jang et al. [35] reported that all of the purified ACE inhibitors from the mushroom *Pleurotus cornucopiae* were noncompetitive inhibitors. Most peptides were competitive inhibitors. Competitive inhibitors have a stronger affinity for ACE than for substrates. They can preferentially bind to ACE active centers and occupy sites such that substrates cannot bind to ACE and are degraded. Peptides with competitive inhibition modes usually bind to the ACE active sites through hydrogen bonding [36, 37] and even coordinate with Zn^2+^ of ACE [38]. Previous studies showed that the amino acids of ACE-inhibitory peptides at the C-terminal bound to ACE. In contrast, the hydrophobic amino acids at the N-terminal interacted with Zn^2+^ ions at the metal-centered active site of ACE or certain active amino acid residues of ACE, thereby inactivating ACE active sites and inhibiting ACE activity [39, 40]. The Lineweaver–Burk plot in the present study showed that the regression curves of pearl matrix peptides with different concentrations intersected at the 1/[*v*] axis, indicating that the peptides were in competitive inhibition mode like most inhibitory peptides.

### 3.3. Activity mechanism of KKCHFWPFPW

Molecular docking is a general method used for predicting the interactions of the inhibitor as a donor and the enzyme as a receptor by calculating the affinity energies, indicating the binding sites and interactive bonds of the donor and the receptor. The molecular docking against ACE was performed by the Maestro 2022-1 Glide module to clarify the ACE-inhibitory mechanism of KKCHFWPFPW.

The structures of the peptide and the peptide–ACE complex are shown in Figure 7, indicating that the peptide was a linear peptide. Figure 8 shows the interaction between the protein and the polypeptide analyzed by Protein-Ligand Interaction Profiler(PLIP), using protein as the reference chain for interaction analysis, with gold representing the protein and blue representing the polypeptide. As shown in Figure 8A, hydrogen bonds were formed between PH-8 of the polypeptide and THR-302 of the protein (solid blue line) between TRP-10 of the polypeptide and GLU-376 of the protein and between TRP-10 of the polypeptide and THR-301 and LYS-449 of the protein. As shown in Figure 8B, both hydrogen bonds and salt bridges were formed between LYS-1 of the polypeptide and GLU-342 of the protein (yellow dashed line). Also, hydrogen bonds and salt bridges were formed between HIS-4 of the polypeptide and ASP-164 and ASN-167 of the protein. As shown in Figure 8C, hydrogen bonds were formed between LYS-2 of the polypeptide and LEU-306 of the protein. In addition, several sets of hydrophobic interactions were present (gray dashed line).

**Figure 7.**
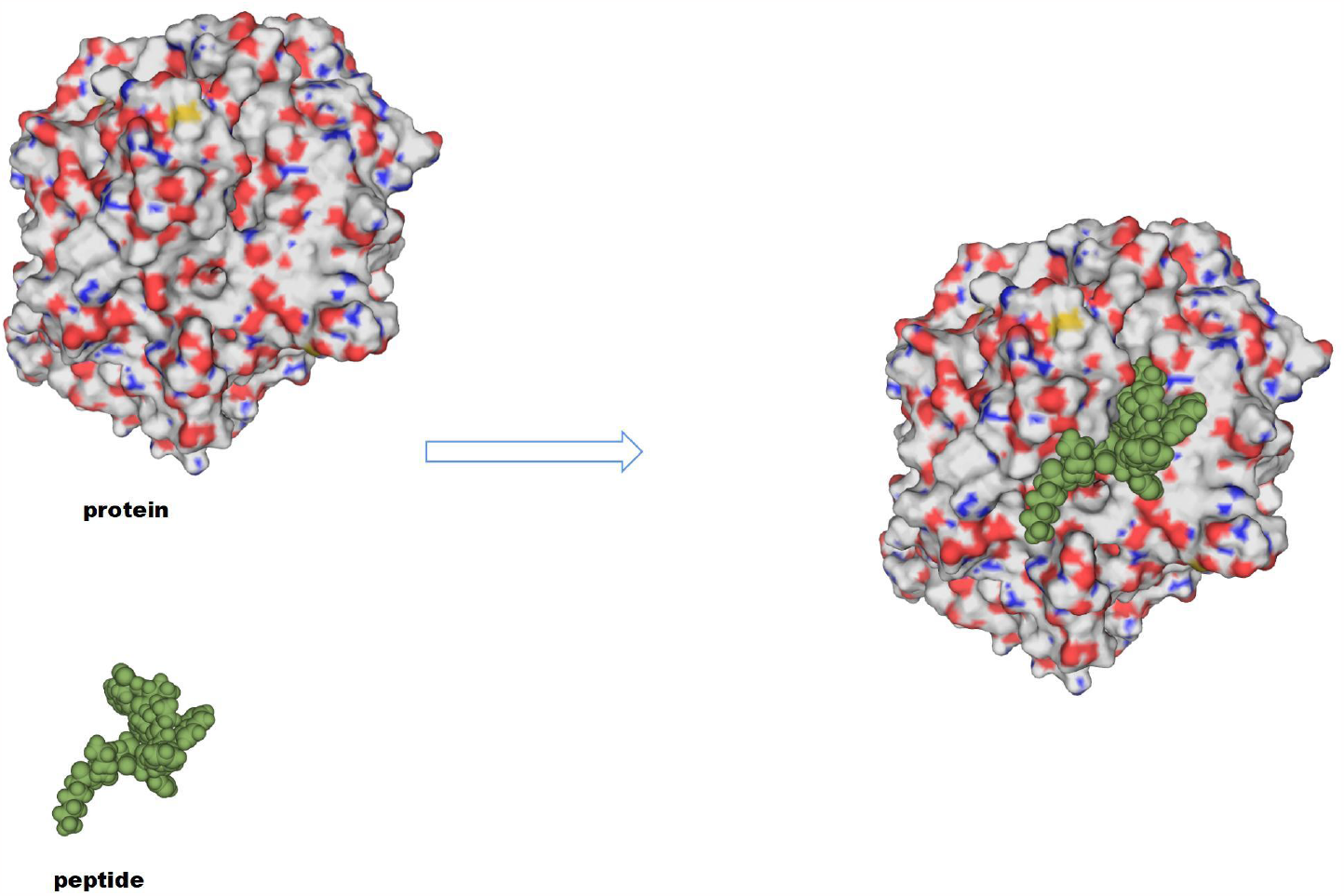
Structures of the peptide and the peptide–ACE complex.

**Figure 8.**
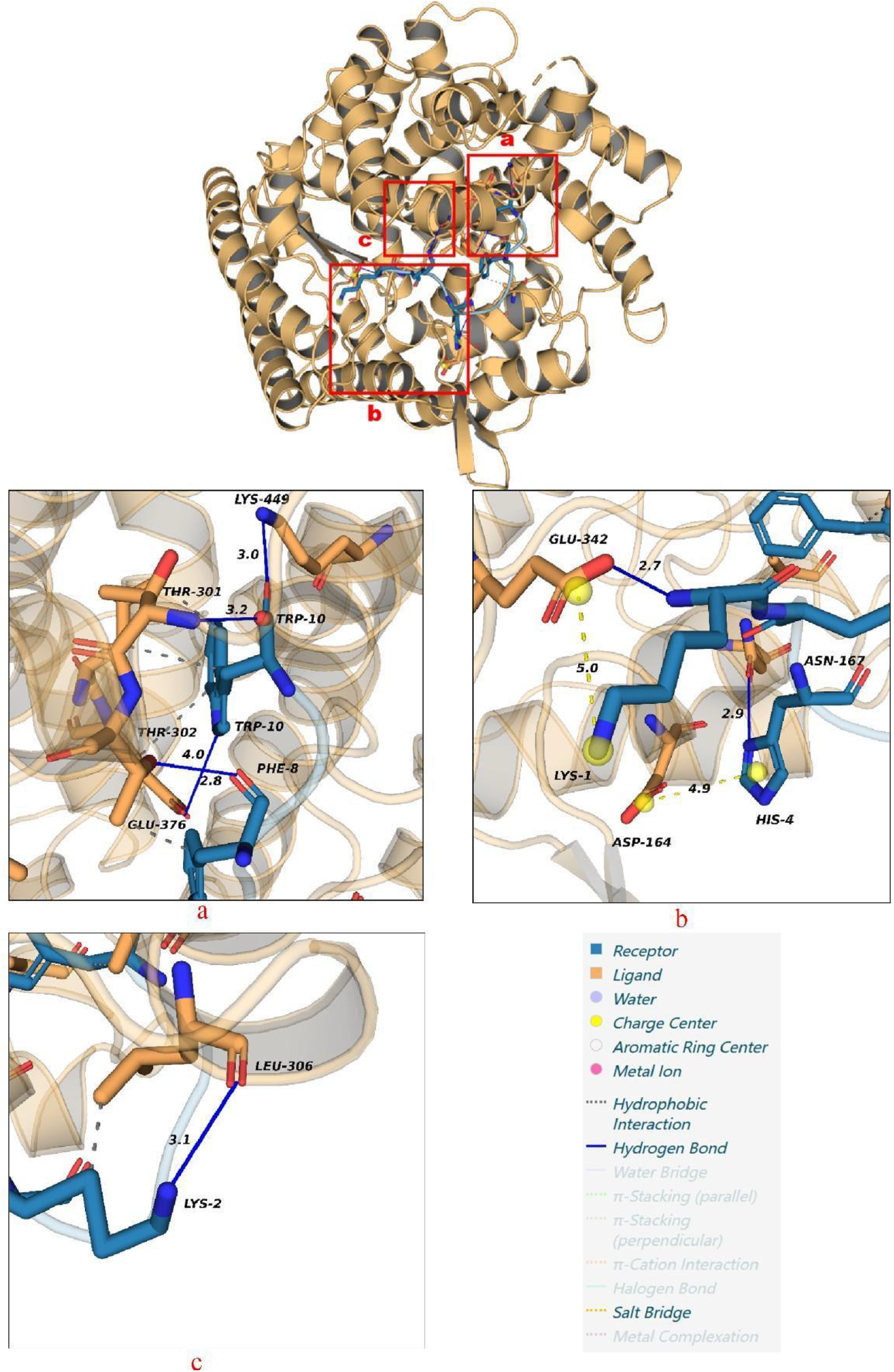
The interaction of KKCHFWPFPW against ACE(PDB: 1O8A) analyzed by PLIP.

## 4. Conclusions

The novel ACE-inhibitory peptide KKCHFWPFPW is a peptide located in the pearl matrix protein with an IC50 value of 4.17 μM, which is competitively inhibited. Potential ACE inhibition mechanisms are mainly attributed to hydrogen bonding and hydrophobic and electrostatic interactions. The peptide can be used in developing health-care products or drugs for treating hypertension, and has broad application prospects in these fields.

